# EGF-Activated Grb7 Confers to STAT3-Mediated EPHA4 Gene Expression in Regulating Lung Cancer Progression

**DOI:** 10.1101/2020.10.24.353268

**Authors:** Pei-Yu Chu, Yu-Ling Tai, Ming-Yang Wang, Hsinyu Lee, Wen Hung Kuo, Tang-Long Shen

## Abstract

Growth factor receptor bound protein-7 (Grb7) is a multi-domain signaling adaptor protein that regulates various cellular functions acting as an adaptor protein in relaying signal transduction. Although several studies indicated that Grb7 amplifies EGFR-mediated signaling in cancers, the detailed regulatory mechanism of whether and how Grb7 is involved in EGFR-mediated lung cancer progression remains unclear. Here, we demonstrate that EGF-regulated Grb7 phosphorylation promotes lung cancer progression through phosphorylation of STAT3. Intrinsically, EGF/EGFR signal is required for the formation of Grb7/STAT3 complex as well as its nuclear accumulation. Once in the nucleus, STAT3 interacts with *EPHA4* promoter, which in turn affects the gene expression level of *EPHA4* through transcriptional regulation. Functionally, EphA4 together with EGFR promotes cancer migration, proliferation, and anchorage-independent growth. Our study reveals a novel mechanism in which Grb7 contribute to lung cancer malignancies through its interaction with STAT3 that leads to sequential regulation of *EPHA4* gene expression in an EGF/EGFR signal-dependent manner.

## 1. Introduction

Lung cancer is the leading cause of cancer-related mortality for both men and women worldwide^1^. Particularly, non-small cell lung cancer (NSCLC), which is the major type of lung cancer, that still remains difficult to treat due to poorly understood pathological regulatory mechanisms ^2^. According to the high frequencies of *EGFR* amplification and mutation in NSCLC patients, EGFR is emerging as one of the first molecules selected for the development of targeted therapies in NSCLC^3, 4, 5^. However, only in recent years has the decreased efficacy of drug or drug resistance emerged in NSCLC patients with anti-EGFR therapy^6, 7^. Considering the significant role of EGFR in NSCLC, it is urgent to elucidate the detailed regulatory mechanisms underlying the EGFR-mediated signal in NSCLC to improve patient outcome in human lung cancer diseases.

Growth factor receptor bound protein-7 (Grb7) is a non-catalytic adaptor protein that modulates cellular functions via the interaction of specific signaling molecules with the protein domains of Grb7 to transmit signal transduction pathways^8^. Indeed, Grb7 is originally identified as a binding partner to activated EGFR^9^. Moreover, our studies indicate that EGF-induced Grb7 phosphorylation/activation is one of the critical steps in cancer development^10^. The co-expression of Grb7 and EGFR in advanced human cancers, such as esophageal cancer, has been reported clinically^11^. Recent clinical studies have indicated that Grb7 is associated with node-positive breast cancer, brain metastasis, decreased survival, and cancer recurrence (especially in breast cancers), suggesting Grb7 as a valuable prognostic marker and therapeutic target^12, 13^. As our studies and others have illustrated, Grb7 amplifies oncogenic signalings to promote cancer progression and metastasis, thereby highlighting Grb7 as a crucial mediator in cancer development^14, 15, 16^. While, Grb7 is implicated in cancer malignancy, the detailed regulatory mechanism of whether and how EGFR-mediated Grb7 signal promotes cancer development needs further investigation.

The Eph receptor family constitutes the largest group of transmembrane receptor tyrosine kinases^17^. Based upon sequence similarity and their ligand-binding specificities, Eph receptors are subdivided into the A and B subclasses (EphA receptor and EphB receptors, respectively), which preferentially bind to ephrinA and ephrinB ligands, respectively^18, 19^. Compared to other Eph receptors, EphA4 is distinguished by its ability to bind and respond to both ephrinA and most ephrinB ligands ^18^. Numerous studies have indicated that the overexpression of EphA4 often correlates with cancer aggression in colorectal, gastric, pancreatic cancers, and glioma^20, 21, 22, 23^. In a recent study, a positive correlation between increased expression of EphA4 protein and STAT3 transcription factor was observed^24^. Possible explanation is that the promoter region of *EPHA4* gene contain several transcription factor binding sites for STAT3^24^, suggesting a high possibility that active STAT3 transcription factor regulates EphA4 protein expression through transcriptional regulation. In addition, EphA4 regulate cell functions by interacting with growth factor receptors, such as fibroblast growth factor (FGF) receptor or EGFR^25, 26^, suggesting the important effects of trans-activation or the synergistic effect of EphA4 and growth factor receptors in mediating signal transduction pathways and functions. Even if the contribution of EphA4 contributes to cancer development has been indicated, the detailed regulatory mechanisms of EphA4 expression remain elusive. Moreover, the molecular and functional effects of EphA4 on growth factor receptors-mediated cancer progression are not well established.

In the present study, we first identified that EGF/EGFR signal mediates the formation of Grb7/STAT3 complex. Moreover, Grb7 is involves in EGF-regulated STAT3 phosphorylation, and enhances nuclear accumulation of STAT3. Together, Grb7-bound STAT3 subsequently induces *EPHA4* gene expression by interacting with the *EPHA4* promoter. Consequently, we revealed that EphA4 amplifies EGFR-mediated cancer aggressiveness. This is the first time that the mechanism in EGF signaling mediates cancer progression involves Grb7/STAT3 complex nuclear accumulation to regulate EphA4 expression to promote cancer aggressiveness.

## 2. Results

### 2.1. EGF/EGFR signal-stimulated Grb7 phosphorylation in the modulation of NSCLC malignancies

Previously, we showed the pivotal role of EGF/Grb7 signal pathway in breast cancer malignancy^10^. Despite the importance of EGFR signal in the development of human NSCLC, it is poorly understood how important the downstream molecule, Grb7, takes part in EGFR-mediated NSCLC tumorigenesis. Here, we demonstrated that tyrosine phosphorylation of Grb7 is elevated by EGF/EGFR signal in H460, H1299, A549 human NSCLC cell lines and HARA human lung squamous carcinoma cell line (Fig. 1a). Impaired cell proliferation (Fig. 1b), cell migration (Fig. 1c), anchorage-independent growth (Fig. 1d), and cell invasion (Fig. 1e) were observed following the knockdown of Grb7 in A549 cells even under EGF stimulation. The observed phenotypes induced by Grb7 knockdown is rescued by the overexpression of Grb7 but not the overexpression of Grb7 SH2 domain (Fig. 1b, c). Consequently, these evidences lead us to speculated an essential role for Grb7 in regulating the development of EGF-mediated NSCLC malignancies.

**Fig. 1.**
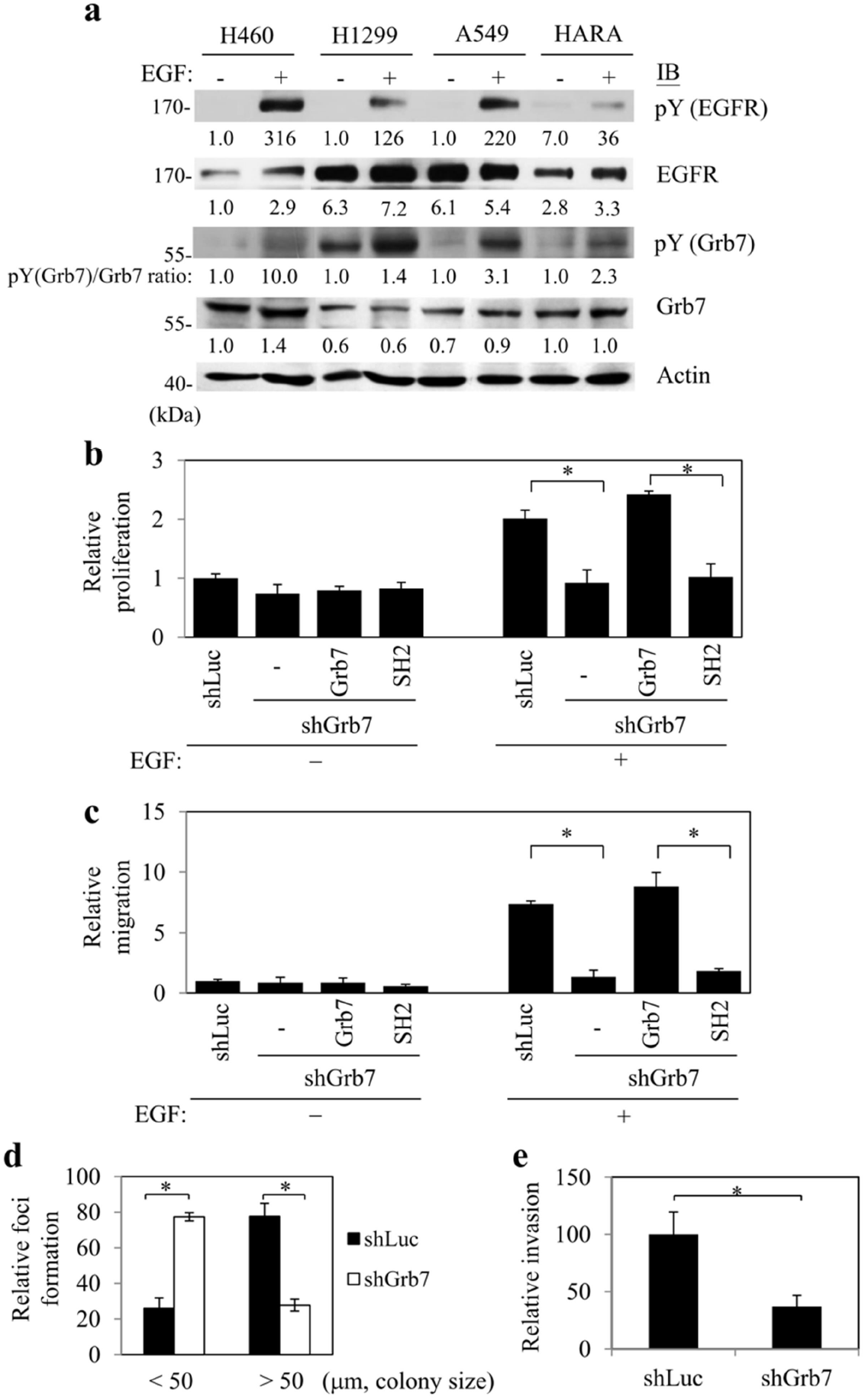
EGF-regulated Grb7 phosphorylation in NSCLC aggressiveness. **a** Cell lysates from serum starved-H460, H1299, A549 human non-small cell lung cancer cell (NSCLC) and HARA human lung squamous carcinoma cell lines with or without EGF (10 ng/ml) treatment were collected and immunoprecipitated with anti-EGFR or anti-Grb7 antibodies followed by Western blot analysis (IB) with an anti-phosphotyrosine antibody to examine the effects of EGF on Grb7 phosphorylation. Cell lysates were also collected and subjected to Western blot with anti-EGFR or anti-Grb7 antibodies. Here, actin was used as a control. A549 cells which were infected with lentiviruses encoding short hairpin Grb7 (shGrb7) were transfected with HA-tagged Grb7 or its truncation mutant, SH2 domain, and subjected to (**b**) cell proliferation assay (for 24 h) by BrdU incorporation and (**c**) cell migration assay (for 8 h) by an modified Boyden chamber assay in the presence of EGF. A549 cells which were infected with lentiviruses encoding short hairpin RNA targeting Grb7 (shGrb7) were subjected to (**d**) anchorage-independent growth assay (for 2 weeks) and (**e**) matrigel invasion assay (for 24 h) in the presence of EGF. HA-overexpressed cells (-) or shLuc-infected (shLuc) cells were used as control cells. All results are shown as the mean ± SEM of at least three independent experiments. Error bars represent ± SEM. * *p* < 0.05.

### 2.2. Grb7 enhances the phosphorylation and the nuclear accumulation of STAT3 in an EGF-dependent manner

In accordance with previous results indicating the indispensable role of EGF-mediated STAT3 activation on tumorigenesis^27^, we investigated whether the status of STAT3 phosphorylation is affected by Grb7 in EGF-stimulated A549 cells (Fig. 2a). Consistently, STAT3 phosphorylation was markedly ablated by the inhibition of EGFR tyrosine kinase activity upon EGF stimulation (Fig. 2a); however, the decrease in the STAT3 phosphorylation was not significantly influenced by Src or FAK (Supplementary Fig. 1). As expected, pharmacological inhibition of STAT3 phosphorylation reduced anchorage-independent growth of NSCLC even in an EGF-dependent manner (Fig. 2b), reinforcing the functional effect of EGF/EGFR signal-mediated STAT3 phosphorylation on NSCLC aggressiveness. Interestingly, overexpression of exogenous Grb7 led to enhanced STAT3 phosphorylation in response to EGF stimulation (Fig. 3a). In contrast, STAT3 phosphorylation was abolished by the knockdown Grb7 even under EGF-stimulated condition (Fig. 3b), suggesting the importance Grb7 to be a crucial mediator in the signal cascade of EGF-induced STAT3 phosphorylation.

**Fig. 2.**
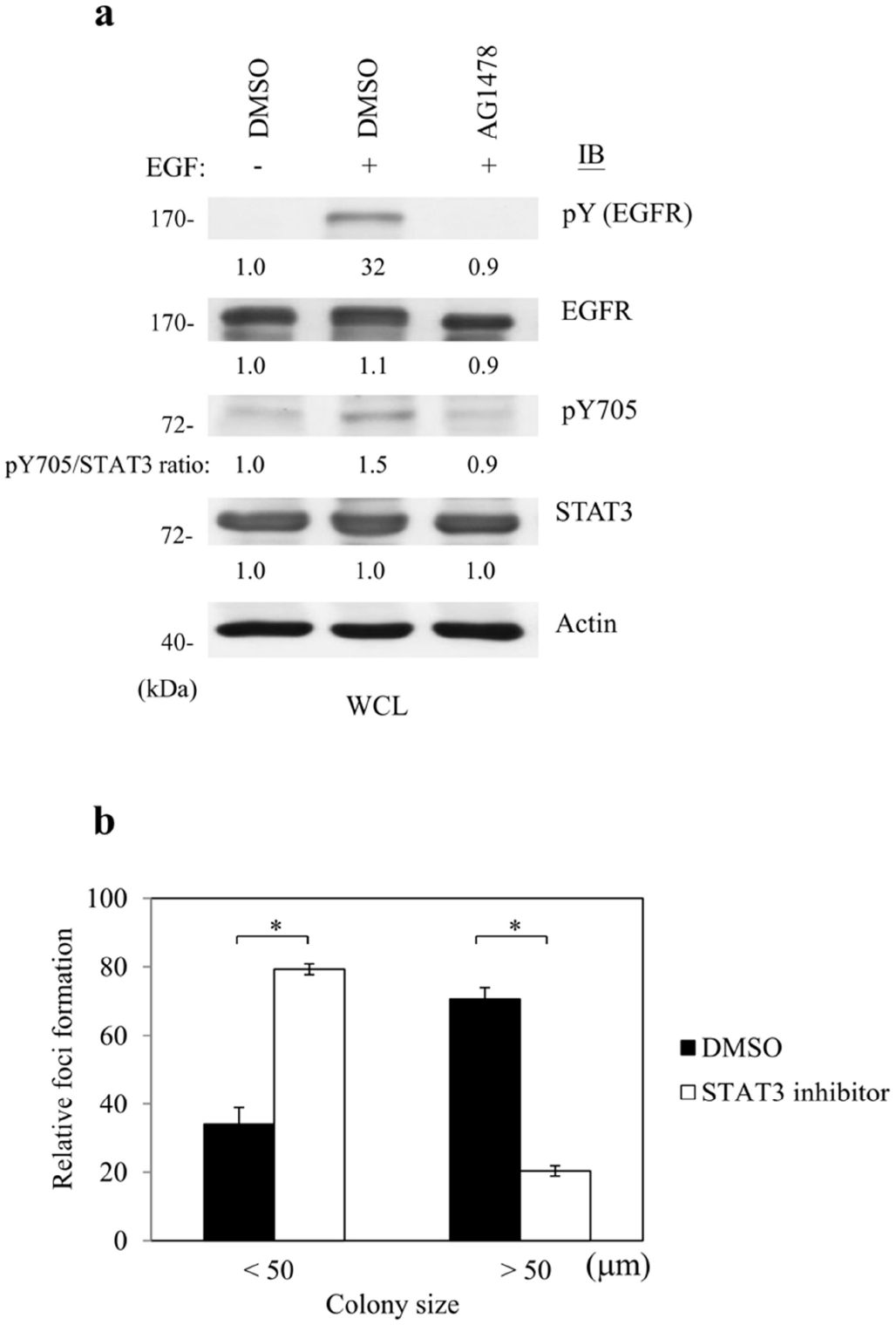
EGF-regulated STAT3 phosphorylation in NSCLC aggressiveness. **a** Serum-starved A549 cells were treated with or without EGF (10 ng/ml) in the presence of EGFR inhibitor AG1478 (5 μM) for 15 min. Cell lysates were collected and subjected to Western blot with anti-EGFR, anti-pTyr1068-EGFR, anti-STAT3, anti-pTyr705-STAT3, and anti-actin antibodies to examine the effects of EGF on protein expression or activity of indicated molecules. **b** A549 cells were treated with STAT3 inhibitor VI (50 μM) and subjected to anchorage-independent growth assay (for 2 weeks) in the presence of EGF. Here, DMSO-treated cells were used as control cells. Results are shown as the mean ± SEM of at least three independent experiments. Error bars represent ± SEM. * *p* < 0.05.

**Fig. 3.**
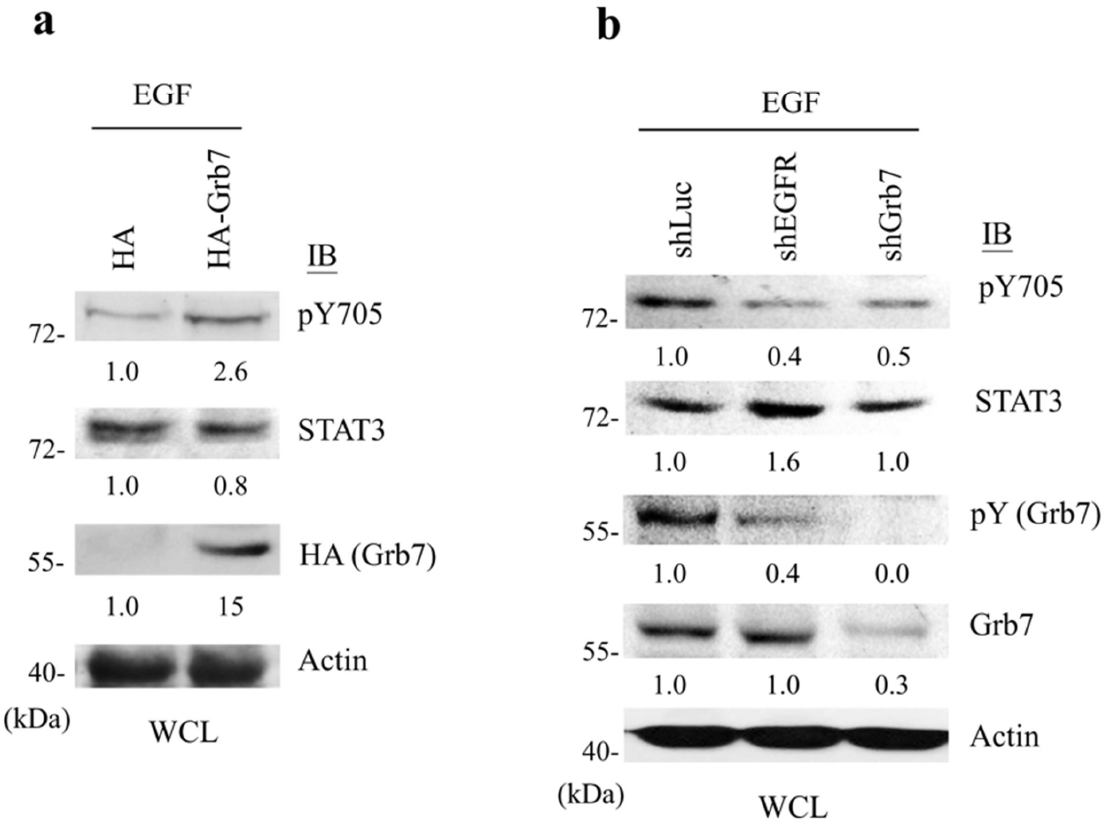
Grb7 involves in STAT3 phosphorylation in an EGF/EGFR signal-dependent manner. **a** A549 cells over-expressing HA-tagged Grb7 were serum-starved for 24 h and then stimulated with EGF (10 ng/ml) for 15 min. Cell lysates were collected and subjected to Western blot with anti-STAT3, anti-pTyr705-STAT3, and anti-actin antibodies to examine the effects of Grb7 on EGF-mediated STAT3 phosphorylation. Grb7 expression was visualized by anti-HA antibody. Here, HA-transfected cells were used as control cells. **b** A549 cells were infected with lentiviruses encoding short hairpin RNA targeting EGFR (shEGFR) or Grb7 (shGrb7) then first serum-starved and stimulated with EGF (10 ng/ml) for 15 min. Cell lysates were collected and immunoprecipitated by an anti-Grb7 antibody followed by Western blotting with an anti-phosphotyrosine antibody. Cell lysates were also collected and subjected to Western blot with anti-STAT3, anti-pTyr705-STAT3 anti-Grb7, and anti-actin antibodies. Here, shLuc-infected A549 cells were used as control cells. WCL, whole cell lysate.

Protein-protein interaction is crucial for intracellular communications. To better understand the mechanistic insight of Grb7 conferring to the EGF-induced STAT3 phosphorylation, the interaction between Grb7 and STAT3 was examined in the EGF-stimulated condition. Upon EGF stimulation, Grb7 could form a complex with STAT3 in the cytosol and the nucleus in a time-dependent manner (Fig. 4a). Interestingly, Grb7 bound to STAT3 was highly phosphorylated (Fig. 4a), implicating the requirement of Grb7 phosphorylation for the physical interaction between STAT3 and Grb7. Moreover, we found that EGF signaling facilitated STAT3 nuclear accumulation in accordance with the increase in the binding ability of Grb7 to STAT3 (Fig. 4b). Given that the phosphorylation of STAT3 at tyrosine 705 is critical for STAT3-mediated cell functions^28^, we further examined its involvement in Grb7-mediated signaling in NSCLC. By employing a dominant-negative point mutant, Y705A, in STAT3, the mutation significantly diminished the interaction with Grb7 in comparison to wild-type (Fig. 4c), indicating the necessity of STAT3 phosphorylation at Y705A for the formation of Grb7/STAT3 complex. Moreover, the Grb7/STAT3 complex and the phosphorylation of STAT3 at tyrosine 705 were impaired in the presence of the Grb7 SH2 domain in a dose-dependent manner compared to the Grb7 RA domain in response to EGF stimulation(is the presence of Sh2 domain suggesting overexpression?) (Supplementary Fig. 2a). In accordance with Fig. 3, our result further suggested that the EGF/EGFR signal mediates the formation of the Grb7/STAT3 complex in concert with the elevated phosphorylation of STAT3. Moreover, the nuclear translocation of both Grb7 and STAT3 was ablated upon blocking the Grb7/STAT3 complex formation through overexpression of Grb7 SH2 domain even in EGF-stimulated condition (Supplementary Fig. 2b). Taken together, these results suggested that the EGF/EGFR signal-stimulated Grb7 phosphorylation is required for the physical interaction and phosphorylation of STAT3, and subsequently the activated status of STAT3, leading to the nuclear translocation of the Grb7/STAT3 complexes in lung cancer cells.

**Fig. 4.**
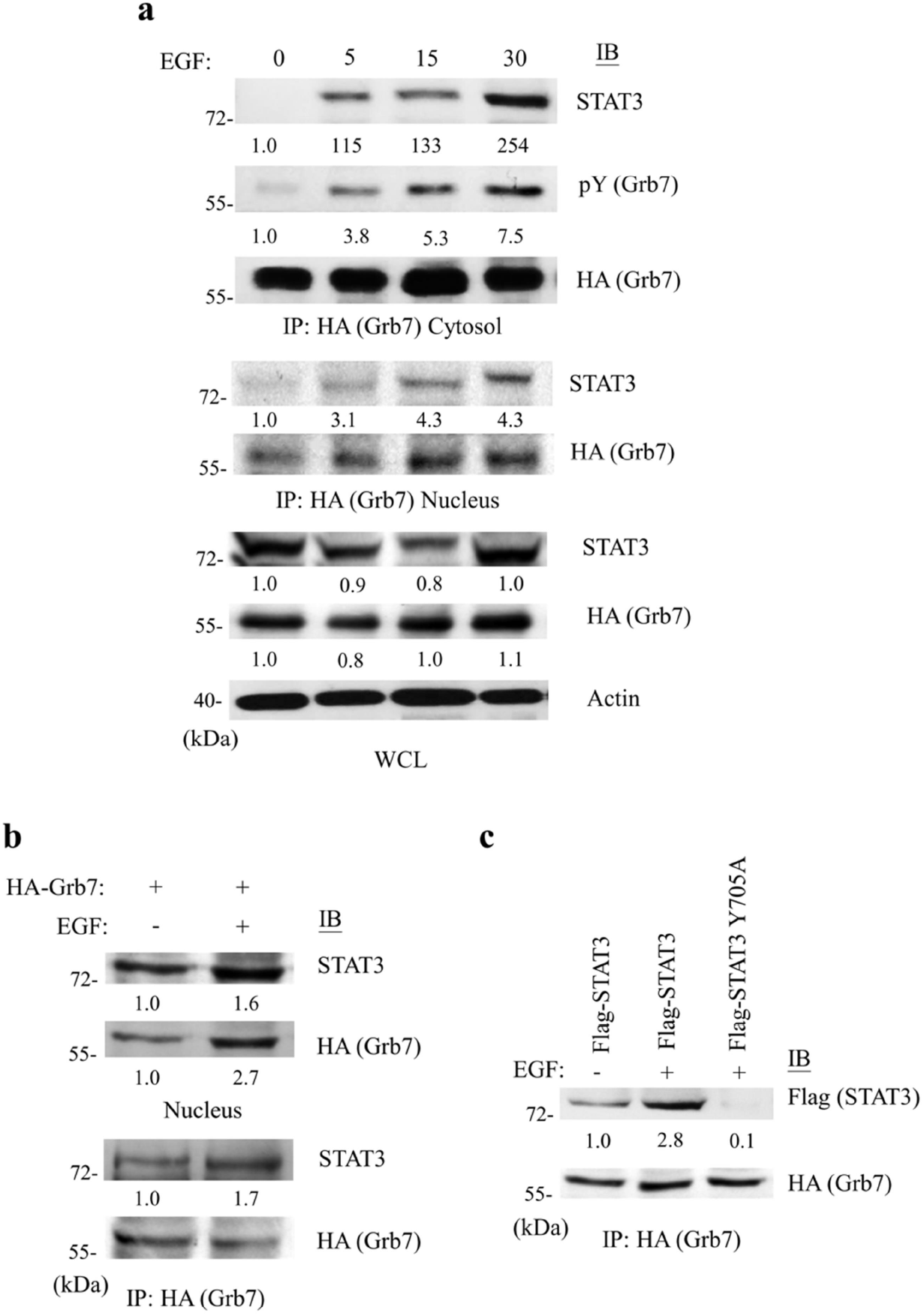
Grb7 interacts with STAT3 and promotes STAT3 nuclear accumulation. **a** HA-tagged Grb7-transfected A549 cells were first serum-starved and then stimulated with EGF (10 ng/ml) for the indicated times (min). Cell lysates were collected and immunoprecipitated by anti-HA antibody against HA-tagged Grb7. The tyrosine phosphorylation of Grb7 and the co-immunoprecipitated STAT3 were visualized by anti-phosphotyrosine and anti-STAT3 antibodies, respectively. Nuclear proteins from A549 cells, described above, were collected and immunoprecipitated by anti-HA antibody against HA-tagged Grb7, and the co-immunoprecipitated STAT3 was visualized by anti-STAT3 antibody. The interaction between STAT3 and Grb7 in response to EGF stimulation was investigated in the cell nucleus. Here, Cell lysates were also collected and subjected to Western blot with anti-STAT3 and anti-actin antibodies. Grb7 expression was visualized by anti-HA antibody. **b** A549 cells over-expressing HA-tagged Grb7 were first serum-starved for 24 h and stimulated with EGF (10 ng/ml) for 15 min. Nuclear proteins from A549 cells were collected and subjected to Western blot with anti-STAT3 antibody and anti-HA antibody against HA-tagged Grb7 to examine effects of Grb7 on the nuclear accumulation of STAT3. Cell lysates were also collected and immunoprecipitated by anti-HA antibody against HA-tagged Grb7, and the co-immunoprecipitated STAT3 was visualized by anti-STAT3 antibody. **c** A549 cells were co-transfected with HA-tagged Grb7 and Flag-tagged STAT3 or its tyrosine to alanine point mutation mutant, Y705A, and cells were first serum-starved and then stimulated with or without EGF (10 ng/ml) for 15 min to investigate the effect of tyrosine phosphorylation of STAT3 on its Grb7-binding ability. Cell lysates were collected and immunoprecipitated by anti-HA antibody against HA-tagged Grb7, and the co-immunoprecipitated STAT3 was visualized by anti-Flag antibody against STAT3.

### 2.3. STAT3 enhances EPHA4 gene expression by interacting with the EPHA4 promoter in an EGF/Grb7 signal-dependent manner

To better understand the molecular mechanism of how nuclear Grb7/STAT3 complex-mediates signal transduction to affect lung cancer, we first evaluated gene expression changes in control and the Grb7-knockdown cells by microarray analysis, which was analyzed by scatter plot of log2 fold-change values of genes. Our results demonstrated that *EPHA4* gene expression level is down-regulated in Grb7-knockdown cells compared to control cells (Supplementary Fig. 3a). Consistently, the EphA4 mRNA level showed a significant decrease in the Grb7-knockdown cells compared to the control cells as shown by Northern blot (Fig. 5a) and quantitative real-time PCR (Supplementary Fig. 3b) analyses. In contrast, the attenuated expression of *EphA4* mRNA (Fig. 5a and Supplementary Fig. 3b) in Grb7-knockdown cells was rescued by overexpression of exogenous Grb7. These results revealed that a novel role of Grb7 is involved in the regulation of *EPHA4* gene expression in lung cancer cells. On the other hand, the *EPHA4* promoter region had been reported to display transcription factor binding sites for STAT3^24^. To further investigate whether the Grb7/STAT3 complex takes part in the expression of *EPHA4* mRNA in the nucleus, chromatin immunoprecipitation (ChIP) assay with anti-STAT3 antibody was performed in A549 human lung cancer cells. In response to EGF/EGFR signal, the interaction between STAT3 and the *EPHA4* promoter region was significantly enhanced in the EGF-stimulated cells compared to the control cells (Fig. 5b).

**Fig. 5.**
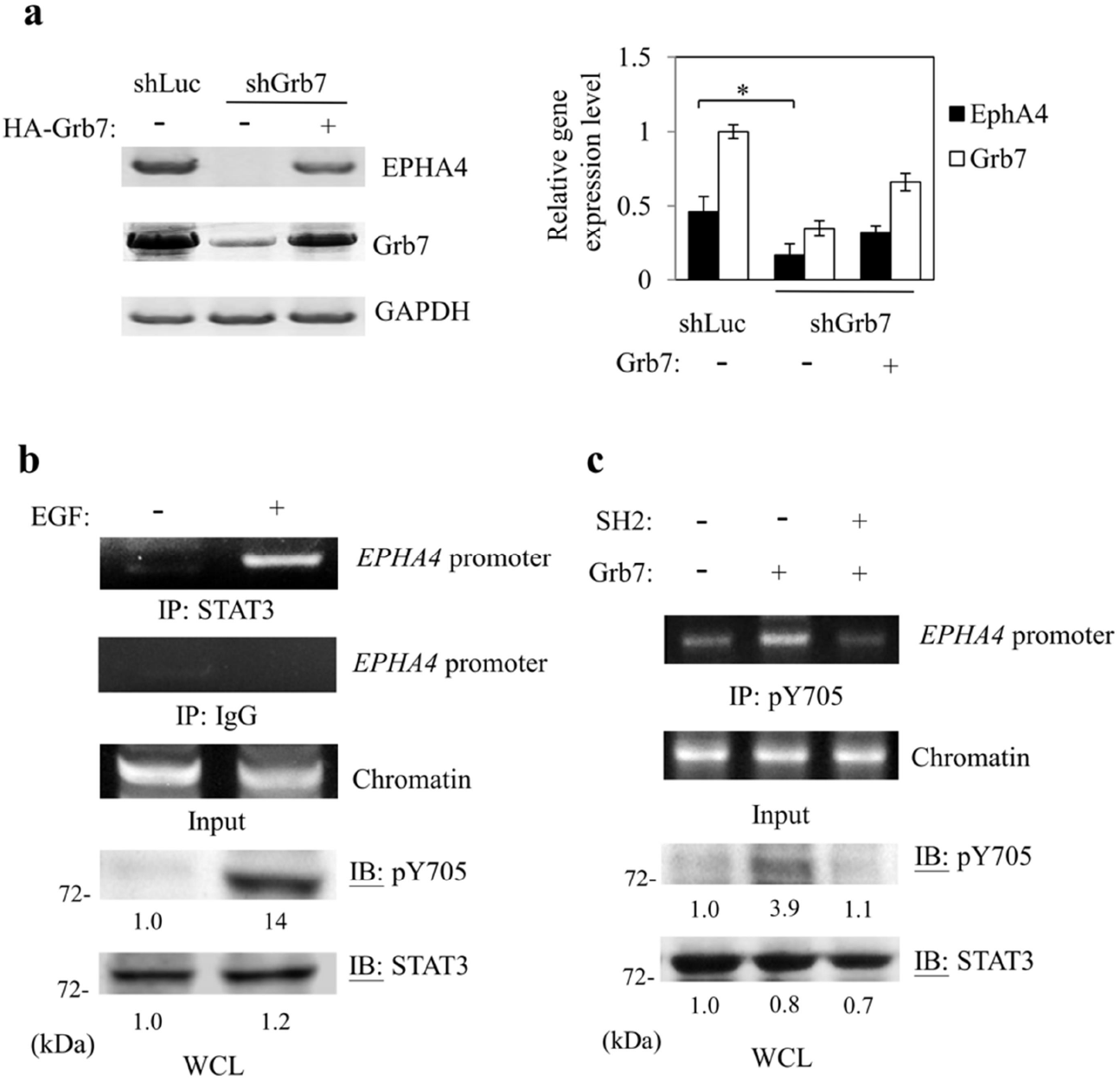
STAT3 interacts with the *EPHA4* promoter in an EGF/Grb7 signal-dependent manner. **a** Grb7 knockdown (shGrb7) A549 cells were transfected with or without exogenous Grb7 (HA-tagged Grb7). mRNA levels of EphA4, Grb7, and GAPDH from A549 cells were detected by Northern blot analysis to investigate effects of Grb7 on the mRNA expression level of EphA4. Here, shLuc-infected A549 cells were used as control cells. GAPDH was used as a control in Northern blot analysis 42. The mRNA levels of Grb7 and EphA4 were normalized to GAPDH (Right). **b** Chromatin immunoprecipitation (IP) assays of the *EPHA4* promoter by STAT3. Soluble chromatin from serum-starved A549 cells stimulated with or without EGF was immunoprecipitated with anti-STAT3 or anti-IgG antibodies. Immunoprecipitates were subjected to PCR with a primer-pair specific to the *EPHA4* promoter to examine the interaction between STAT3 and *EPHA4* promoter. **c** HA-tagged Grb7 was co-transfected with or without Grb7 SH2 domain into A549 cells. Serum-starved A549 cells, described above, were stimulated with EGF (10 ng/ml). Then, soluble chromatin was collected to conduct chromatin immunoprecipitation assays by using an anti-pTyr705-STAT3 antibody and followed by PCR analysis with a primer-pair specific to the *EPHA4* promoter. Results indicated that Grb7 facilitated interaction between phosphorylated-STAT3 and *EPHA4* promoter in response to EGF stimulation, whereas Grb7 SH2 domain impaired the interaction between phosphorylated-STAT3 and *EPHA4* promoter even in the EGF stimulation.

Furthermore, to examine whether Grb7-mediated phosphorylation at tyrosine 705 of STAT3 participates in the binding of STAT3 to *EPHA4* promoter in an EGF-dependent manner, we performed ChIP-PCR analyses. Our results showed that phosphorylated STAT3 bound *EPHA4* promoter was significantly increased in the Grb7-overexpressed condition compared to the control (Fig. 5c and Supplementary Fig. 4); whereas, a strong repression of this interaction was observed in cells overexpressing Grb7-SH2 domain (Fig. 5c). Similarly, the dominant-negative point mutant, Y705A, of STAT3 led to an absence of *EPHA4* promoter binding ability even under EGF stimulation or/and overexpression of exogenous Grb7 (Supplementary Fig. 4).

Next, we investigated the transcriptional activity of the STAT3/Grb7 complex on the *EPHA4* promoter using a transcriptional reporter assay. Upon EGF stimulation, *EPHA4* promoter activity was significantly increased in STAT3-overexpressed cells (Fig. 6a). On the contrary, the *EPHA4* promoter activity was dramatically reduced by exogenous overexpression of a dominant-negative point mutant, Y705A, of STAT3 even in an EGF-stimulated condition (Fig. 6a). Consistently, a decrease in the *EPHA4* promoter activity was also observed in the Grb7-knockdown cells compared to the control cells, whereas the luciferase activity was rescued by exogenous addition of Grb7 (Fig. 6b). To highlight the above observation, we further dissected the Grb7/STAT3 complex response element within the *EPHA4* promoter region, the promoter activity of *EPHA4* gene with or without Grb7 expression was first investigated by promoter deletion analysis. Here, we generated a series of *EPHA4* promoter truncation mutants (Supplementary Fig. 5a) and subjected them to the luciferase reporter assay. As shown in Supplementary Fig. 5b, luciferase activity showed a significant difference between an overexpressed Grb7 and an endogenous Grb7 in *EPHA4* promoter −1954/+91 bp and −1001/+91 bp. Moreover, deletion in the region from −1001 to −248 bp diminished the effect of Grb7 on its contribution towards *EPHA4* promoter activity (Supplementary Fig. 5b). Consistent with the effect of Grb7 on *EPHA4* promoter activity, a significant effect of STAT3 on *EPHA4* promoter activity was detected in *EPHA4* promoter −1954/+91 bp and −1001/+91 bp (Supplementary Fig. 5c). As expected, the EphA4 protein expression level is down-regulated in the STAT3-knockdown cells (Fig. 6c) and the Grb7-knockdown cells (Fig. 6d) compared to control. Similarly, the attenuated EphA4 protein expression level in Grb7-knockdown cells was rescued by overexpression of exogenous Grb7 (Fig. 6d). Together, these results indicate that the *EPHA4* promoter region contained the Grb7 and STAT3 response elements and EGF/EGFR signal is involved in the expression of *EPHA4* gene in lung cancer mediated by Grb7/STAT3.

**Fig. 6.**
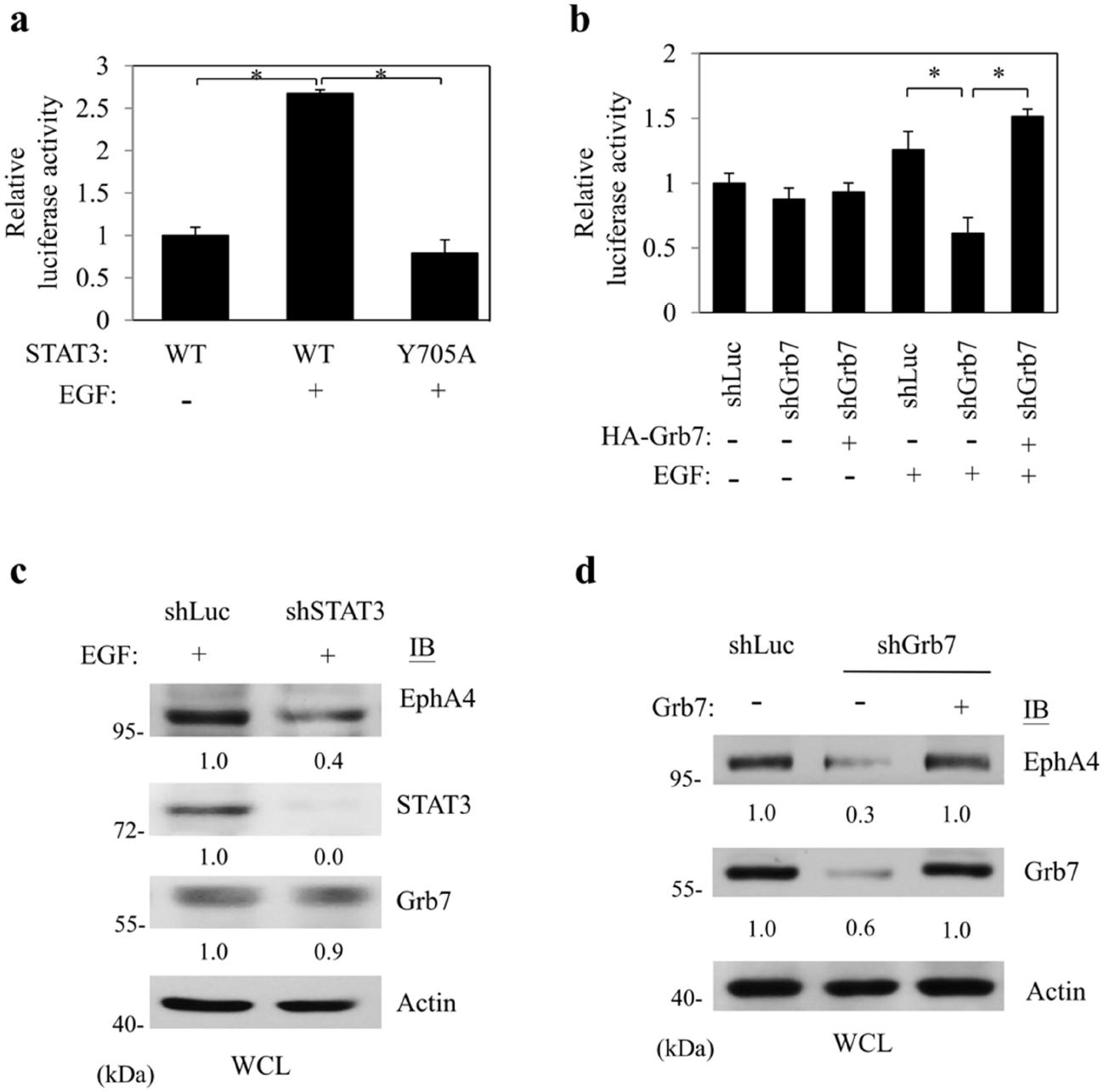
EGF/Grb7 signal induces the binding of STAT3 to *EPHA4* promoter and the facilitating EphA4 protein expression. **a** The −1954 to +91 construct of *EPHA4* promoter region was co-transfected with Flag-tagged STAT3 or its tyrosine to phenylalanine mutant, Y705F, into A549 cells. Thirty-six hours post-transfection, cells were stimulated with or without EGF (10 ng/ml) and assayed for luciferase activity. Results are shown as the mean ± SEM of at least three independent experiments. Error bars represent ± SEM. * *p* < 0.05. **b** The −1954 to +91 construct of *EPHA4* promoter region was co-transfected with or without HA-tagged Grb7 into Grb7 knockdown (shGrb7) A549 cells. Thirty-six hours post-transfection, cells were first serum starved and stimulated with EGF (10 ng/ml) and assayed for luciferase activity. Here, HA-overexpressed cells were used as control cells. All results are shown as the mean ± SEM of at least three independent experiments. Error bars represent ± SEM. * *p* < 0.05. **c** STAT3 knockdown (shSTAT3) A549 cells were first serum starved and stimulated with EGF (10 ng/ml). Cell lysates were collected and subjected to Western blot with anti-STAT3, anti-EphA4, anti-Grb7, or anti-actin antibodies to investigate effects of STAT3 on protein expression levels of EphA4. Here, shLuc-infected cells were used as control cells. **d** shGrb7-infected A549 cells were transfected with or without HA-tagged Grb7. Thirty-six hours post-transfection, cells were first serum starved and stimulated with EGF (10 ng/ml). Cell lysates were collected and subjected to Western blot with anti-EphA4, anti-Grb7, or anti-actin antibodies to invesitgate effects of Grb7 on protein expression levels of EphA4. Here, actin was used as a control. HA-overexpressed cells or shLuc-infected cells were used as control cells.

### 2.4. EphA4 interacts with EGFR and amplifies EGFR-mediated cancer proliferation and anchorage-independent growth

Previous studies indicated that the synergistic response of EphA4 and growth factor receptors is involved in cancer progression^22, 29^. As a crucial oncoprotein in lung cancers, we investigated whether up-regulation of EphA4 synergize with EGFR signaling to promote lung cancer malignancy *in vitro.* Indeed, upon EGF/ephrin A1 (EFNA1) co-stimulated condition, prominent effects on cancer cell proliferation (Fig. 7a) and anchorage-independent growth ability (Fig. 7b) were seen.

**Fig. 7.**
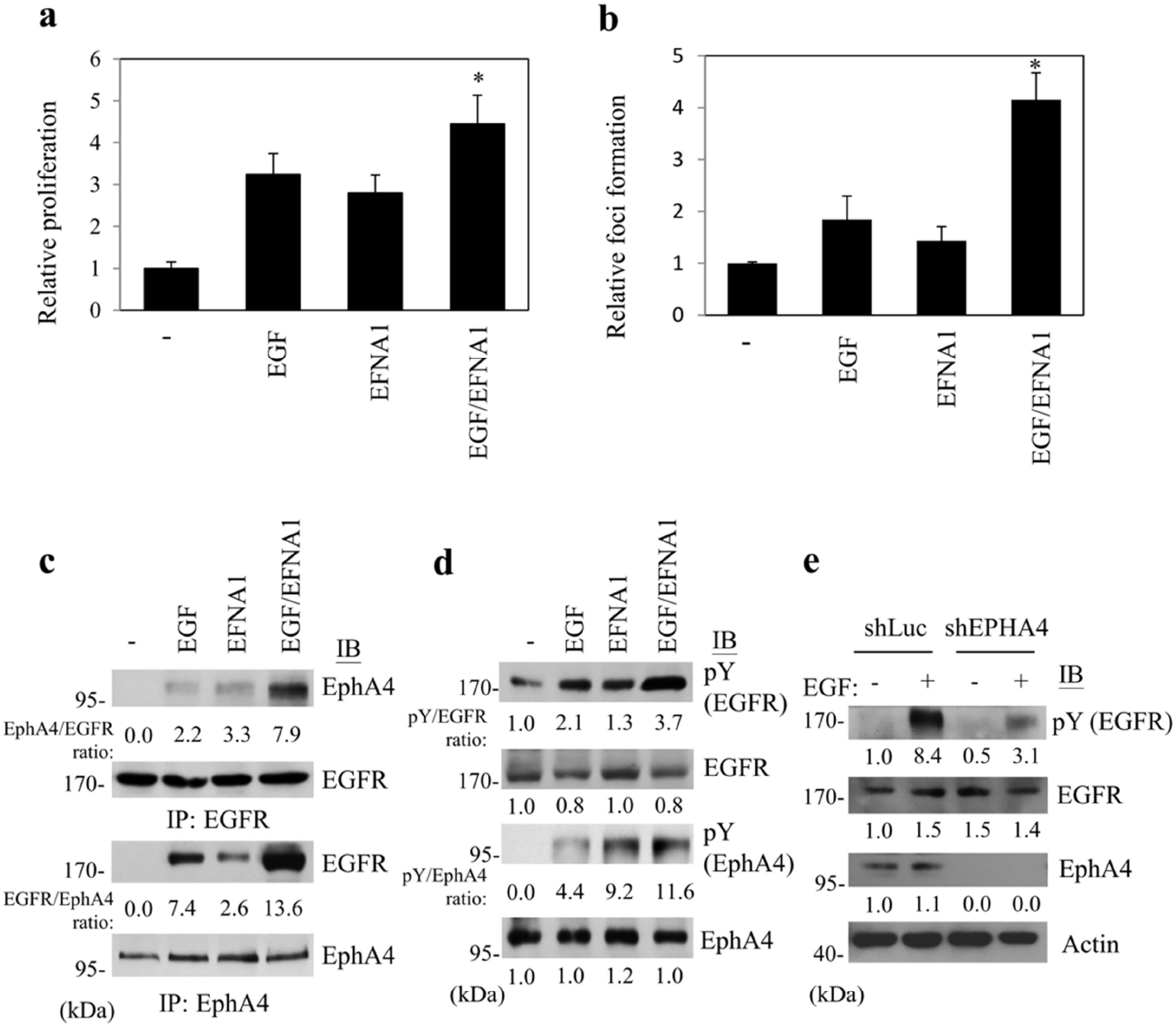
EphA4 interacts with EGFR and amplifies EGFR-mediated cell functions. **a, b** A549 cells were subjected to cell proliferation assay (for 24 h) by (**a**) BrdU incorporation analysis and (**b**) soft agar assay (for 2 weeks) to investigate anchorage-independent growth ability in the presence of EGF, EFNA1, or both EGF and EFNA1 (EGF/EFNA1) to examine effects of the activated EphA4 and EGFR on cell functions. All results are shown as the mean ± SEM of at least three independent experiments. Error bars represent ± SEM. * *p* < 0.05. **c** Cell lysates from serum-starved A549 cells stimulated with EGF (10 ng/ml), EFNA1 (0.5 μg/ml), or both EGF and EFNA1 were collected and immunoprecipitated by anti-EGFR or anti-EphA4 antibodies, and the co-immunoprecipitated EphA4 or EGFR, were visualized by anti-EphA4 and EGFR antibodies, respectively, to examine the interaction between EGFR and EphA4. Here, the immunoprecipitated EGFR or EphA4 was subjected to Western blot with anti-phosphotyrosine antibody. **d** Results indicated tyrosine phosphorylation of EGFR was the highest in both EGF and EFNA1 co-stimulated condition. **e** Cell lysates from EPHA4 knockdown (shEPHA4) A549 cells stimulated with or without EGF (10 ng/ml) were collected and immunoprecipitated by an anti-EGFR antibody, and the immunoprecipitated EGFR was subjected to Western blot with anti-phosphotyrosine antibody to examine effects of EphA4 on EGFR tyrosine phosphorylation. The protein expression of EGFR, EphA4, or actin was detected by anti-EGFR, anti-EphA4, or anti-actin antibodies, respectively.

In light of the above findings, we further investigated whether EphA4 binding to EGFR occurs in lung cancer cells. Although the EGFR/EphA4 complex is formed upon only EGF- or EFNA1-stimulated condition, the strong interaction between EGFR and EphA4 was found in the EGF and EFNA1 co-stimulated condition (Fig. 7c). Moreover, tyrosine phosphorylation of EphA4 or EGFR was increased in response to EGF or EFNA1 stimulation, respectively (Fig. 7d), suggesting the mutual transactivation between EGFR and EphA4. Not surprisingly, tyrosine phosphorylation of both EGFR and EphA4 was strongly increased in EGF and EFNA1 co-stimulated condition (Fig. 7d) and tyrosine phosphorylation of EGFR was decreased in the EphA4 knockdown cells compared to control cells even under the EGF-stimulated condition (Fig. 7e). Collectively, our findings suggest that EphA4 interacts with EGFR, resulting in the amplification of EGFR-mediated NSCLC aggressiveness.

### 2.5. EphA4 amplifies EGFR-mediated ERK1/2 phosphorylation

To reveal the downstream signaling of the activated EGFR/EphA4 complex, we first examined the phosphorylation of ERK1/2, an important downstream of EGFR/Grb7 signal transduction^2^ in response to EGF/EFNA1 co-stimulation. As shown in Fig. 8a, a profound impact on enhanced phosphorylation on ERK1/2 was observed in the EGF and EFNA1 co-stimulated condition. Consistently, the MEK inhibitor ablated the activated EGFR/EphA4 complex-mediated ERK1/2 phosphorylation (Fig. 8b). Functionally, the MEK inhibitor also prevented EGF/EFNA1-triggered anchorage-independent growth ability of lung cancer cells (Fig. 8c). These results indicated that ERK1/2 is involved in the activated EGFR/EphA4 complex-regulated NSCLC aggressiveness.

**Fig. 8.**
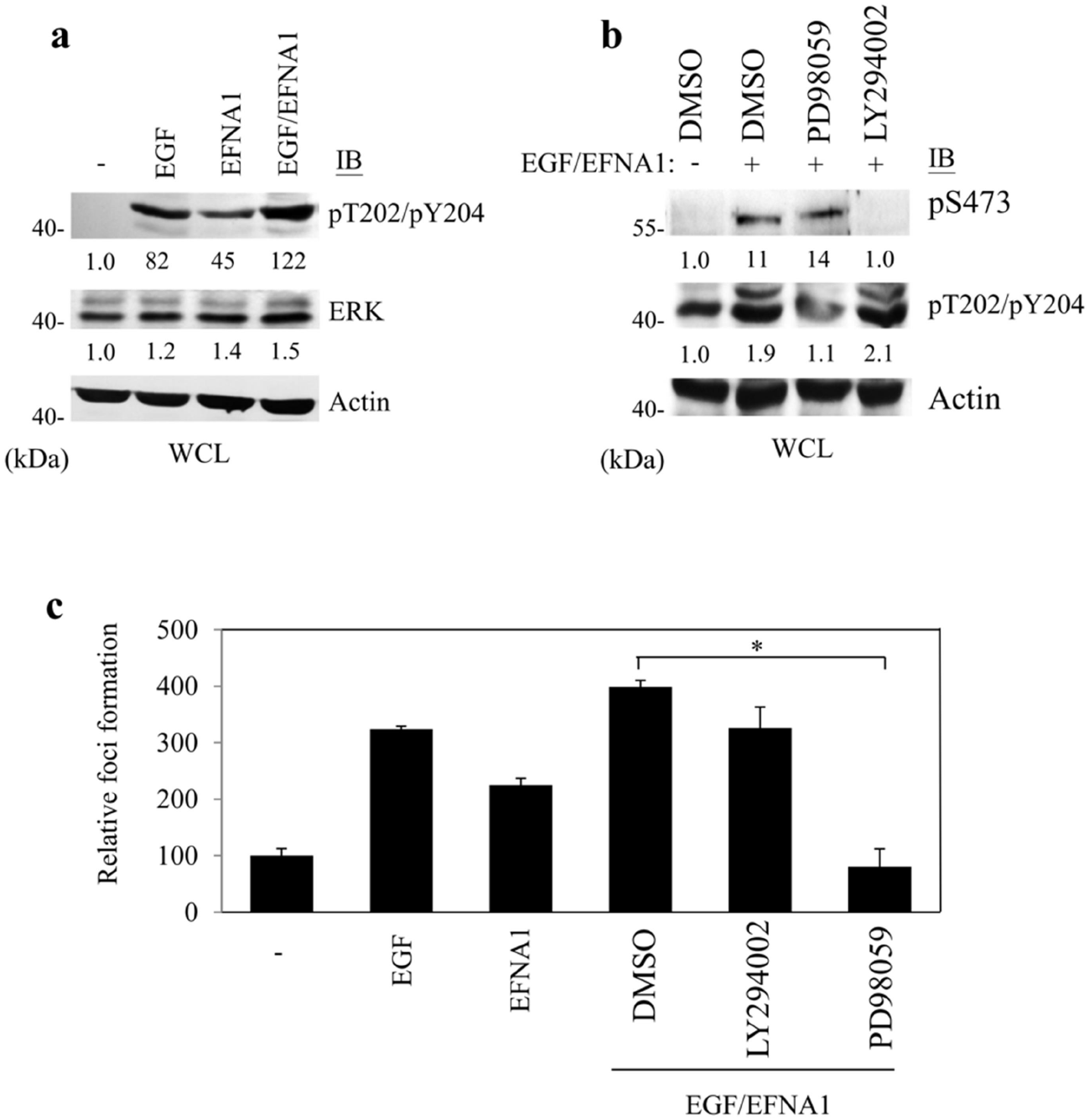
EphA4 amplifies EGFR-mediated ERK1/2 phosphorylation in the regulation of anchorage-independet growth. **a** Cell lysates from serum-starved A549 cells stimulated with EGF (10 ng/ml), EFNA1 (0.5 μg/ml), or both EGF and EFNA1 (EGF/EFNA1) were collected and subjected to Western blot with anti-ERK or anti-pT202/pY204-ERK antibodies to examine effects of EphA4 on EGFR-mediated ERK phosphorylation. **b** Cell lysates from MEK inhibitor PD98059-or PI3K inhibitor LY294002-treated A549 cells stimulated with both EGF and EFNA1 were collected and subjected to Western blot with anti-S473-AKT or anti-pT202/pY204-ERK antibodies. **c** MEK inhibitor PD98059- or PI3K inhibitor LY294002-treated A549 cells were subjected to examine the anchorage-independent growth ability (for 2 weeks) in the presence of both EGF and EFNA1. Here, DMSO, EGF, or EFNA1 treatments were used as controls. Results are shown as the mean ± SEM of at least three independent experiments. Error bars represent ± SEM. * *p* < 0.05.

### 2.6. Expression of Grb7 and EphA4 positively correlated with lung cancer aggressiveness

To assess the relationship of EphA4 as well as Grb7 expression pattern and the pathophysiological role in lung adenocarcinoma, we analyzed the protein expression of EphA4 and Grb7 in established human lung carcinoma cell lines (CL1-0, CL1-1, CL1-2, and CL1-5) that display progressive aggressiveness. Consistent with an aggressive role for the EphA4 and Grb7 in lung cancer (Fig. 1b, 1c, 1d, 1e, 7a and 7b), increased protein expression pattern of EphA4 as well as Grb7 was found to be markedly associated with the aggressiveness of lung cancers (Supplementary Fig. 6).

## 3. Discussion

Grb7 functions as a critical mediator during cancer development, whereas the regulatory mechanism of Grb7 in EGF/EGFR signal-mediated cancer aggressiveness is not well established. Here, our results reveal, for the first time, a regulatory process of EGF/EGFR-mediated Grb7 signal in NSCLC aggressiveness. We found that Grb7 involves in the phosphorylation and the nuclear accumulation of STAT3 in response to EGF/EGFR signal. Subsequently, STAT3 mediates the expression of *EPHA4* gene through binding to *EPHA4* promoter in a Grb7-dependent manner. As a result, EphA4 protein binds to EGFR and amplifies EGFR-mediated NSCLC aggressiveness. Our results highlight a novel regulatory mechanism of Grb7 by the regulation of STAT3-modulated *EPHA4* gene expression in EGF/EGFR signal-mediated NSCLC development (Fig. 9).

**Fig. 9.**
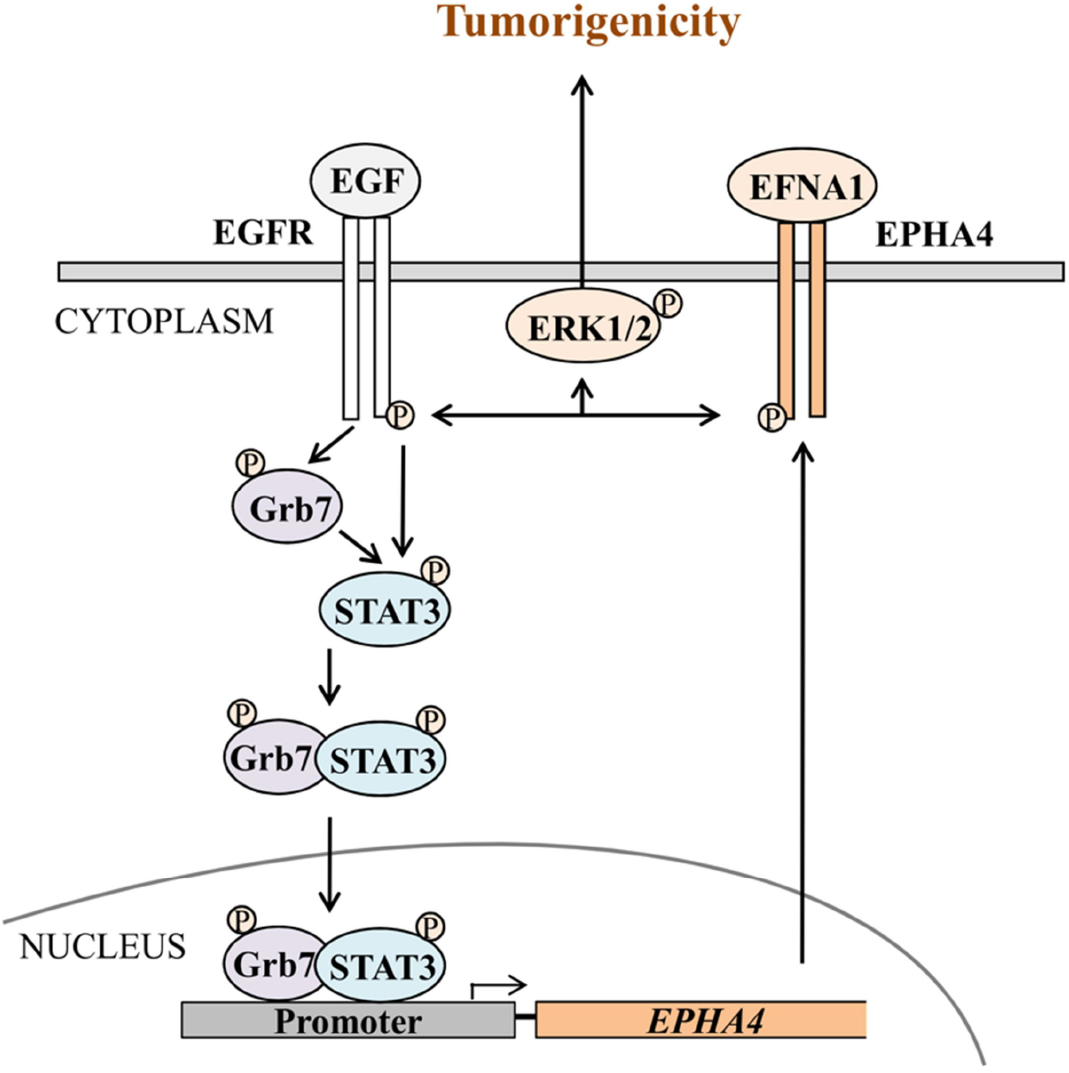
The working model for the EGF/Grb7/STAT3 signal-mediated NSCLC malignancy via the regulation of EphA4 expression.

Tumor microenvironmental factors exhibit significant effects on cancer development^30^. In line with previous studies^27^, our studies indicated that EGF stimulates STAT3 phosphorylation as well as STAT3-mediated cancer malignancy (Fig. 2a, b). Importantly, we further identified that Grb7 is required for EGF-mediated STAT3 activation (Fig. 3) and cancer malignancy (Fig. 1b,c). In response to the upstream EGR/EGFR signal activation, we found that Grb7 interacts with the phosphorylated STAT3 (Supplementary Fig. 2a); whereas, defective STAT3 phosphorylation significantly ablated the Grb7 binding ability of STAT3 (Fig. 4c). Additionally, EGF/EGFR signal induced the interaction between Grb7 and STAT3 in concert with elevated phosphorylation of STAT3 (Fig. 3,Fig. 4, and Supplementary Fig. 2a). These studies suggested that the phosphorylation status of STAT3 determines its association with Grb7 and Grb7-bound STAT3 might stabilize the activated status of STAT3 in response to the stimulation of tumor microenvironmental cues.

Consistent with the functions of Grb7 as an adaptor protein, the subcellular localization of Grb7 can be investigated not only in the cytoplasm^10, 14^ but also in the nucleus^31, 32^. With respect to these findings, our studies indicated that Grb7 can be detected in both cytoplasm and nucleus (Fig. 4a). Although signals and functional effects of cytoplasmic Grb7 on cancer malignancy is illustrated in many studies ^10, 14, 15, 16^, little is known about the pathological role of nuclear Grb7 during cancer development. Our studies first found that increased nuclear Grb7 can be detected upon EGF signal treatment compared to the one without EGF stimulation (Fig. 4b). At the same time, the interaction between Grb7 and STAT3 is strongly detected in the nucleus (Fig. 4a). Nuclear localization of Grb7 has been shown to be facilitated by its calmodulin-binding domain that displays a sequence of high similarity to the nuclear localization signal^31^. Thus, we hypothesize that Grb7 may function as a carrier protein in the regulation of the nuclear transport of STAT3. Nevertheless, the detailed regulatory mechanism will be required to illustrate the proposed model of the Grb7-mediated STAT3 shuttling between the nucleus and the cytoplasm.

Due to the functional impacts of transcription factor-mediated reprogramming on the regulation of cancer malignancy ^33^, our studies provide a novel evidence for STAT3 transcription factor-mediated *EPHA4* gene expression through transcriptional regulation in a Grb7-dependent manner (Fig. 5 and Supplementary Fig. 5). The overexpression of EphA4 or up-regulation of EphA4-mediated signals are often correlated with malignancy, increased tumor relapse, or drug resistance in many kinds of cancers including breast cancer, glioma, lung cancer, or multiple myeloma ^22, 34, 35, 36^. Clinically, overexpression of EphA4 was consistently investigated in lung cancer samples compared to normal samples^36^. In line of previous studies^36^, our results indicated significant up-regulation of EphA4 as well as Grb7 in concert with the progressive invasiveness of human lung carcinoma (Supplementary Fig. 6), highlighting the functional effects of Grb7 on the EphA4-mediated lung cancer malignancy through STAT3-induced *EPHA4* gene transcriptional program.

The crosstalk between membrane receptors results in signal cascades amplifying synergistic effects during cancer development^37, 38^. Indeed, we found the strong interaction between EphA4 and EGFR in response to the both EGF and EFNA1 stimulation (Fig. 7c). Simultaneously, the phosphorylation of EphA4 and EGFR as well as downstream ERK 1/2 signaling is increased upon the co-stimulation of EGF and EFNA1 (Fig. 7a, Fig. 8a). Functionally, a significant increase in the malignancy of NSCLC can be investigated in response to the co-stimulation of EGF and EFNA1 (Fig. 7a,b); whereas, the inhibition of ERK 1/2 ablated the functional effects of the co-stimulation of EGF and EFNA1 (Fig. 8c).

In conclusion, our studies illustrated that EGF/EGFR-Grb7 signal regulates STAT3-induced *EPHA4* gene expression in lung cancer aggression. Subsequently, EphA4 interacts with EGFR and amplifies EGFR-mediated cancer malignancy. The significant increase of downstream ERK 1/2 signal functions as the result of EGFR and EphA4 signal cascade-amplifying synergistic effects. Our results demonstrate a novel regulatory mechanism involved in the modulation of malignancy in NSCLC.

## 4. Methods

### 4.1. Reagents and antibodies

Protein A-Sepharose 4B, 2-Morpholin-4-yl-8-phenylchromen-4-one (LY294002, PI3K inhibitor), N-(3-chlorophenyl)-6,7-dimethoxy-4-quinazolinamine (AG1478, EGFR inhibitor), ephrin-A1/Fc fragment, human IgG/Fc fragment, 5’-bromo-2-deoxyuridine (BrdU), and the mouse monoclonal anti-BrdU antibody were purchased from Sigma-Aldrich (St Louis, MO). Herceptin (Trastuzumab) was obtained from Roche Applied Science (South San Francisco, CA). The 2-(2-amino-3-methoxyphenyl)-4H-1-benzopyran-4-one (PD98059, MEK inhibitor) and STAT3 inhibitor VI were from Calbiochem (Darmstadt, Germany). The EGF was from Millipore (Billerica, MA). The Dual-Luciferase Reporter Assay System was from Promega (Madison, WI). Lipofectamine 2000^TM^, Opti-MEM, Dulbecco’s modified Eagle’s medium (DMEM), and RPMI-1640 were obtained from Invitrogen (Carlsbad, CA). The mouse monoclonal anti-phosphotyrosine (PY99), anti-HA (12CA5), anti-GFP (B-2) and the rabbit polyclonal anti-Grb7 (C-20), anti-EGFR (1005), and anti-EphA4 (S-20) antibodies were obtained from Santa Cruz Biotechnology (Santa Cruz, CA). The mouse monoclonal anti-phospho-EGFR (Tyr-1068), and anti-actin (C4) antibodies were obtained from Millipore (Billerica, MA). The rabbit polyclonal anti-ERK1/2, anti-phospho-ERK (Thr-202/Tyr-204), anti-p38 MAPK, anti-phospho-p38 MAPK (Thr-180/Tyr-182), anti-SAPK/JNK, anti-phospho-SAPK/JNK (Thr-183/Tyr-185), anti-AKT, anti-phospho-AKT (Ser-473), anti-STAT3, and anti-phospho-STAT3 (Tyr-705) antibodies were purchased from Cell Signaling (Danvers, MA).

### 4.2. Cell cultures, transient transfection and lentiviral infection

A549 human non-small cell lung cancer (NSCLC) and 293T human kidney epithelial cell lines were maintained in Dulbecco’s modified Eagle’s medium (DMEM) supplemented with 10% fetal bovine serum (FBS). NCI-H460 (H460) and NCI-H1299 (H1299) human NSCLC cell lines and HARA human lung squamous carcinoma cell line were maintained in RPMI-1640 supplemented with 10% FBS (Gibco BRL, Gaithersburg, MD). Cell transfection was carried out by Lipofectamine 2000^TM^ (Invitrogen) according to the manufacturer’s instructions and as described^14^.

Lentiviruses encoding GRB7, STAT3, or Luciferase small-hairpin RNA (shLuc) were obtained from the TRC lentiviral shRNA library in National RNAi Core Facility of Academia Sinica, Taiwan. The targeting sequencings of various shRNAs were as follows: GRB7 shRNA (clone ID: TRCN0000061387) 5’-CCAGGGCTTTGTCCTCTCTTT-3’; STAT3 shRNA (clone ID: TRCN0000020843) 5’, GCAAAGAATCACATGCCACTT-3’. Production and infection of lentiviruses were processed according to the guidelines of the National RNAi Core Facility of Academia Sinica (Taipei, Taiwan) and also see^14^.

### 4.3. Quantitative real-time PCR and RT-PCR

The cDNA was synthesized from 1 μg of purified RNA derived from A549 cells using M-MLV reverse transcriptase according to manufacturer’s recommendations (Invitrogen, Carlsbad, CA). The housekeeping gene *GAPDH* was used as a reference for normalization (primers: 5’-ACGACCCCTTCATTGACCTC-3’ and 5’-CTTTCCAGAGGGGCCATCCAC-3’). Quantitation of PCR products was estimated by SYBR Green reagent (2X Maxima SYBR Green qPCR Master Mix; Fermentas, Waltham, MA) using the LightCycler^®^ 480 System (Roche, South San Francisco, CA), and the data was analyzed with LightCycler^®^ 480 Gene Scanning Software according to manufacturer’s instructions (Roche, South San Francisco, CA). The qPCRs were performed in quadruplicate, and copy number alterations were scored as validated if 2^ΔCt^ (relative copy number) was ≥2^39^ or ≤0.5 (loss) with CV ≤15% of mean 2^ΔCt^. RT-PCRs were conducted in two-step reactions by using two-step RT-PCR: M-MLV reverse transcriptase in M-MLV RT-PCR system (Promega, Madison, WI) and Platinum *Taq* polymerase (GeneMark, Taipei, Taiwan) sequentially. The *EPHA4* and *GRB7* gene expression levels were normalized by *GAPDH*, which was used as reverse transcription control.

### 4.4. Northern blotting

Total RNA was isolated using TRIzol reagent (Invitrogen, Carlsbad, CA). 20 μg total RNA was electrophoresed through a 1% agarose gel with EtBr. The gel was treated with 0.05N NaOH for 20 min and 7% formaldehyde for another 20 min. The RNA was then transferred onto Nylon membrane (GE Healthcare) by the capillary method and UV cross-linked (1200 × 100 J). Membranes were pre-hybridized for 1 h at 65°C in a Church buffer containing 0.5 M NaHPO4 (pH 7.2), 7% SDS, 1% BSA, and 1 mM EDTA. A 451 bp DNA probe was prepared between bases 564-1014 of *GRB7* (primers: 5’-ACTTCGCCAAGGAAGAACTGTT-3’ and 5’-AACACACGGACT-3’) and bases 348-581 of *EPHA4* (primers: 5’-GACTTGCAAGGAGACGTT-3’ and 5’-AACACACGGACTGATACC-3’). These cDNAs were labeled with [α-^32^P] dCTP (PerkinElmer Inc., Waltham, MA) using the Amersham Rediprime II DNA Labeling System (Amersham Pharmacia, Amersham, UK) according to the manufacturer’s instructions. The membrane was hybridized in Church buffer for 14 h at 65°C in a rotating oven and was washed twice for 10 min each in 2X SSC (20X SSC: 3 M NaCl, 0.3 M sodium citrate, pH 7.0) at room temperature, then in 2X SSC with 1% SDS for 15 min at 65°C, followed by two 15 min high-stringency washes in 0.1% SSC, 0.1% SDS at 65°C. The membrane was exposed to autoradiography film (Kodak, Wilmington, DE) for 2 days at −80°C and developed.

### 4.5. Immunoprecipitation, co-immunoprecipitation and Western blotting analyses

Proteins were extracted and subjected to co-immunoprecipitation and/or Western blotting as described^10^. Briefly, cells were washed twice with ice-cold PBS and then lysed with 1% Nonidet P-40 lysis buffer (20 mM Tris, pH 8.0, 137 mM NaCl, 1% Nonidet P-40, 10% glycerol, 1 mM Na3VO4, 1 mM phenylmethylsulfonyl fluoride, 10 mg/ml aprotinin, and 20 mg/ml leupeptin), harvested by scraping, and on the ice for 30 min. Cell lysates were collected and clarified by centrifugation for 25 min at 4 °C, and total protein concentration was determined using the Bradford Assay according to the manufacturer’s instructions (Sigma-Aldrich, St Louis, MO). Immunoprecipitations were carried out by incubating cell lysates with antibodies as indicated for 12 h at 4 °C, followed by incubation for 4 h with protein A-Sepharose 4B beads. After washing five times with lysis buffer, immune complexes were resolved using SDS-PAGE. Western blotting was proceeded, and then incubated with indicated primary antibodies. The membranes were incubated with horseradish peroxidase-conjugated IgG as a secondary antibody and the Western Lightning^®^-ECL system (PerkinElmer Inc., Waltham, MA) for detection.

### 4.6. Nuclear and cytosolic fractionation

The preparation of nuclear and cytosolic fractions was modified from the procedure described by Chou *et al*.^40^. Briefly, A549 cells were pelleted and washed twice with cold PBS, suspended in 400 μl ice-cold hypotonic lysis buffer (10 mM Hepes pH 7.5, 10 mM NaCl, 2 mM MgCl_2_, 10mM NaF, 1 mM EDTA, 1 mM DTT and 0.1mM Na_3_Vo_4_, supplemented with protease inhibitors). 0.5% NP40 was added to cell mixtures on ice for additional 3 min and then followed with gentle taps to break down cellular membrane. The supernatant was collected as cytoplasmic extract after a centrifugation at 800 *g* for 5 min at 4°C. The pellet containing the nuclei was washed with 1 ml of washing buffer (20-mM Tris-HCl pH 7.9, 140-mM KCl, and 20% glycerol) and then resuspended in 150 μl nucleus extraction buffer (25 mM HEPES pH 7.5, 500 mM NaCl, 5 mM MgCl_2_, 10 mM NaF, 1 mM EDTA, 1 mM DTT, 1% glycerol and 0.2% NP40) for 20 min on ice, centrifuged at 12,000 × *g* for 15 min at 4°C, and the supernatants were collected as nuclear extracts. The same volumes of nuclear or cytosolic fractions were analyzed by immunoprecipitation and Western blotting.

### 4.7. Construction of luciferase reporter gene constructs and promoter luciferase assay

−1954/+91 (from −1954 to +91 bp), −1001/+91 (from −1001 to +91 bp), or −248/+91 (from −284 to +91 bp) of *EPHA4* promoter region were cloned into pGL3-Basic vector (Promega, Madison, WI) to allow transcription of firefly luciferase gene under the control of this fragment. For promoter luciferase assay, cells were co-transfected with luciferase-containing constructs (pGL3) and phRL-TK synthetic renilla vector (Promega, Madison, WI) in a molar ratio of 1:60 (phRL-TK versus pGL3). The level of firefly luciferase activity was normalized to that of *Renilla reniformis* luciferase activity for each transfection. For all experiments, cells were cultured for 48 h after transfection, and cell lysates were prepared and examined by using the Dual luciferase reporter assay system (Promega, Madison, WI), according to the manufacturer’s protocol.

### 4.8. Chromatin immunoprecipitation (ChIP) assay

Chromatin derived from 5 × 10^6^ A549 cells was used for each immunoprecipitation. Cells were cross-linked with 1% formaldehyde for 15 min at room temperature, and were stopped by the addition of 0.125 M glycine. The cells were collected by centrifugation and rinsed in cold PBS. The cell pellets were resuspended in PBS with protease inhibitors (leupeptin and aprotinin, both 100 ng/ml), incubated on ice for 20 min. The nuclei were collected by centrifugation and then resuspended in ChIP sonication buffer (1% Triton X-100, 0.1% deoxycholate, 5 mM EDTA, 50 mM Tris-HCl [pH 8.1], 150 mM NaCl, and protease inhibitors) and incubated on ice for 10 min. Chromatin was sonicated and collected by centrifugation. The samples were immunoprecipitated with anti-STAT3, anti-phospho-STAT3 (Tyr-705), or anti-Grb7 antibody. After overnight incubation at 4°C, add protein A-Sepharose 4B beads to each immunoprecipitation and incubate at 4 °C for 2 h. The immunoprecipitates were then washed twice with 10 ml washing buffer (1% Triton X-100, 0.1% deoxycholate, 150 mM NaCl, 5 mM EDTA, and 10 mM Tris-pH 8.1) and once with LiCl Immune complex wash buffer (0.25 M LiCl, 0.5% deoxycholate, 0.5% NP-40, 1 mM EDTA, 10 mM Tris-pH 8.1). Immunocomplexes were eluted from the beads by adding 250 μl Elution buffer (1% SDS, 0.1 M NaHCO3) followed by incubation at room temperature for 20 min. Protein-DNA cross-links were reversed in 0.25 M NaCl at 65°C for 4 h, after which DNA was isolated by adding ethanol to each sample and placing at −20°C overnight. The samples were resuspended in proteinase K buffer and incubated at 55°C for 1 h. DNAs were extracted with phenol-chloroform-isoamyl alcohol (25:24:1) followed by extraction with chloroform-isoamyl alcohol and then precipitated with 1/10 volume of 3 M NaOAc (pH 5.3), and 2.5 volumes of ethanol. The pellets were collected by centrifugation. DNAs were then resuspended in 100 μl of TE buffer (10mM Tris, 8.1, 1 mM EDTA) and amplified by PCR. Primer pairs for EPHA4 promoter region which contains STAT3 binding sites were identified by PCR using the following primers: region from −1001 to −58 forward 5’-GCTTCCCAGTCCCGGTCT-3’, reverse 5’, AGTTAGGAGAGCAGCGGGCT-3’; region from −502 to −58 forward 5’-CAGGAACAAGGGCCTCTGTCT-3’-reverse 5’-TGTCCCTCTGACAATGTGCCATC-3’.

### 4.9. Boyden chamber assay

Cell migration assays were carried out using a Neuro Probe (Cabin John, MD) 48-well chemotaxis Boyden chamber as described previously^14^. Approximately 1 × 10^5^ cells were resuspended in DMEM and added in the upper wells of the Boyden chamber, and the cells were allowed to migrate toward EGF (10 ng/ml) in DMEM as the chemoattractant or DMEM only as a control in the lower wells for 8 hours in a 37°C humidified incubator. At the end of experiments, cells on the upper side of the polycarbonate membrane were removed and the bottom-side cells were fixed with methanol for 10 minutes and stained with crystal violet (Sigma-Aldrich, St Louis, MO). The migrated cells were counted from five randomly selected fields of each sample under a light microscope (model IX71, Olympus, Japan).

### 4.10. Scattering onto immobilized ligands assay

The movement of A549 cells onto ephrin-A1 was performed as described previously^41^ with modification. Briefly, ephrin-A1/Fc or IgG/Fc was coated on 24-well culture plates at 4°C. The subconfluent A549 cells were plated on coverslips in 24-well plate. After reaching confluence, cells were serum-starved for 24 h. The coverslips with cells were transferred cell-face-up onto the coated 24-well plate containing control medium or medium supplemented with 10 ng/ml EGF. After 48 h, cells were fixed with 4 % paraformaldehyde for 20 min and stained with crystal violet. Coverslips were removed, and the scattered rings of cells were photographed.

### 4.11. BrdU incorporation assay

Cell proliferation was estimated by using BrdU incorporation assay as described previously^14^. After serum starvation for 24 h in DMEM with 0.2% FBS, the subconfluent cells were replated on ephrin-A1/Fc or IgG/Fc for 1 hr, further with or without co-stimulated by EGF for 15 min and then incubated for 24 h in DMEM containing 10% FBS and 100 μM BrdU. Cells were then fixed in 4% paraformaldehyde for 15 min at room temperature and were subjected to immunofluorescent staining. Cells were then counted in 5 fields and scored for BrdU-positive staining in each independent experiment.

### 4.12. Soft agar assay

Anchorage-independent growth was examined using soft agar assays as described previously^14^. We supplemented the cells twice a week with DMEM containing EGF (10 ng/ml), ephrin-A1 (0.5 μg/ml), combination, or STAT3 inhibitor VI (50 μM). After the 2nd week of treatment, colony numbers were scored under a light microscope (model IX71, Olympus, Japan).

### 4.13. Matrigel invasion assay

Cell invasion was analyzed by BD BioCoat™ growth factor reduced Matrigel™ invasion chambers according to the manufacturer’s instructions (BD Biosciences, Mississauga, Ontario, Canada). 5 × 10^4^ serum-starved A549 cells with or without shRNA lentiviruses, as indicated, were used for Matrigel invasion assay. Here, EGF (10 ng/ml) in DMEM was used as a chemoattractant. After incubation for 24 h at 37°C, noninvasive cells were removed and invasive cells were fixed with methanol for 15 min and stained with crystal violet for 15-20 min. The number of invasive cells was counted from three randomly selected fields under a light microscope (model IX71, Olympus, Japan).

### 4.14. Statistical analysis

All results represent the Mean ± SEM of at least three independent experiments. The error bars (SEM) shown in Fig. were derived from biological replicates, not technical replicates. Significant differences between two groups were evaluated using a two-tailed Student’s *t*-test based on the analysis of variance. Statistical difference was considered significant at *p* < 0.05 and indicated in Fig. as *.

## Acknowledgements

This work was supported by the Ministry of Science and Technology, Taiwan (105-2320-B-002-058-MY3) and the Dragon-Gate Program, Ministry of Science and Technology, Taiwan (106-2911-I-002-569).

## Author contributions

P.Y.C. and T.L.S. designed experiments. P.Y.C. and Y.L.T. performed experiments. P.Y.C., Y.L.T., and T.L.S. analyzed the data. M.Y.W., H.L., and W.H.K. contributed to the data discussion. P.Y.C., Y.L.T., and T.L.S. wrote the manuscript. All authors read and approved the final manuscript.

## Competing interests

The authors declare that they have no conflict of interest.

## Additional information

Supplementary information is available.

## References

1. Siegel RL, Miller KD, Jemal A. Cancer statistics, 2019. CA: a cancer journal for clinicians 69, 7–34 (2019).

2. Bethune G, Bethune D, Ridgway N, Xu Z. Epidermal growth factor receptor (EGFR) in lung cancer: an overview and update. Journal of thoracic disease 2, 48–51 (2010).

3. Hynes NE, Lane HA. ERBB receptors and cancer: the complexity of targeted inhibitors. Nature reviews Cancer 5, 341–354 (2005).

4. Janne PA, Engelman JA, Johnson BE. Epidermal growth factor receptor mutations in non-small-cell lung cancer: implications for treatment and tumor biology. Journal of clinical oncology: official journal of the American Society of Clinical Oncology 23, 3227–3234 (2005).

5. Minna JD, Peyton MJ, Gazdar AF. Gefitinib versus cetuximab in lung cancer: round one. Journal of the National Cancer Institute 97, 1168–1169 (2005).

6. Perez-Soler R, et al. Determinants of tumor response and survival with erlotinib in patients with non--small-cell lung cancer. Journal of clinical oncology: official journal of the American Society of Clinical Oncology 22, 3238–3247 (2004).

7. Rho JK, et al. Epithelial to mesenchymal transition derived from repeated exposure to gefitinib determines the sensitivity to EGFR inhibitors in A549, a non-small cell lung cancer cell line. Lung Cancer 63, 219–226 (2009).

8. Shen TL, Guan JL. Grb7 in intracellular signaling and its role in cell regulation. Front Biosci 9, 192–200 (2004).

9. Margolis B, et al. High-efficiency expression/cloning of epidermal growth factor-receptor-binding proteins with Src homology 2 domains. Proceedings of the National Academy of Sciences of the United States of America 89, 8894–8898 (1992).

10. Chu PY, Li TK, Ding ST, Lai IR, Shen TL. EGF-induced Grb7 recruits and promotes Ras activity essential for the tumorigenicity of Sk-Br3 breast cancer cells. J Biol Chem 285, 29279–29285 (2010).

11. Tanaka S, et al. Coexpression of Grb7 with epidermal growth factor receptor or Her2/erbB2 in human advanced esophageal carcinoma. Cancer Res 57, 28–31 (1997).

12. Ramsey B, et al. GRB7 protein over-expression and clinical outcome in breast cancer. Breast cancer research and treatment 127, 659–669 (2011).

13. Nadler Y, Gonzalez AM, Camp RL, Rimm DL, Kluger HM, Kluger Y. Growth factor receptor-bound protein-7 (Grb7) as a prognostic marker and therapeutic target in breast cancer. Annals of oncology: official journal of the European Society for Medical Oncology / ESMO 21, 466–473 (2010).

14. Chu PY, et al. Tyrosine phosphorylation of growth factor receptor-bound protein-7 by focal adhesion kinase in the regulation of cell migration, proliferation, and tumorigenesis. The Journal of Biological Chemistry 284, 20215–20226 (2009).

15. Pradip D, Bouzyk M, Dey N, Leyland-Jones B. Dissecting GRB7-mediated signals for proliferation and migration in HER2 overexpressing breast tumor cells: GTP-ase rules. American journal of cancer research 3, 173–195 (2013).

16. Tai YL, et al. Grb7 Protein Stability Modulated by Pin1 in Association with Cell Cycle Progression. PLoS One 11, e0163617 (2016).

17. Davis S, et al. Ligands for EPH-related receptor tyrosine kinases that require membrane attachment or clustering for activity. Science 266, 816–819 (1994).

18. Kullander K, Klein R. Mechanisms and functions of Eph and ephrin signalling. Nature reviews 3, 475–486 (2002).

19. Pasquale EB. Eph receptor signalling casts a wide net on cell behaviour. Nature reviews 6, 462–475 (2005).

20. Iiizumi M, et al. EphA4 receptor, overexpressed in pancreatic ductal adenocarcinoma, promotes cancer cell growth. Cancer science 97, 1211–1216 (2006).

21. Oshima T, et al. Overexpression of EphA4 gene and reduced expression of EphB2 gene correlates with liver metastasis in colorectal cancer. International journal of oncology 33, 573–577 (2008).

22. Fukai J, Yokote H, Yamanaka R, Arao T, Nishio K, Itakura T. EphA4 promotes cell proliferation and migration through a novel EphA4-FGFR1 signaling pathway in the human glioma U251 cell line. Mol Cancer Ther 7, 2768–2778 (2008).

23. Oki M, Yamamoto H, Taniguchi H, Adachi Y, Imai K, Shinomura Y. Overexpression of the receptor tyrosine kinase EphA4 in human gastric cancers. World journal of gastroenterology: WJG 14, 5650–5656 (2008).

24. Hameetman L, et al. EPHA4 is overexpressed but not functionally active in Sezary syndrome. Oncotarget 6, 31868–31876 (2015).

25. Sawada T, et al. Ternary complex formation of EphA4, FGFR and FRS2alpha plays an important role in the proliferation of embryonic neural stem/progenitor cells. Genes to cells: devoted to molecular & cellular mechanisms 15, 297–311 (2010).

26. Yokote H, et al. Trans-activation of EphA4 and FGF receptors mediated by direct interactions between their cytoplasmic domains. Proceedings of the National Academy of Sciences of the United States of America 102, 18866–18871 (2005).

27. Song L, Turkson J, Karras JG, Jove R, Haura EB. Activation of Stat3 by receptor tyrosine kinases and cytokines regulates survival in human non-small cell carcinoma cells. Oncogene 22, 4150–4165 (2003).

28. Darnell JE, Jr. STATs and gene regulation. Science 277, 1630–1635 (1997).

29. Arvanitis D, Davy A. Eph/ephrin signaling: networks. Genes & Development 22, 416–429 (2008).

30. De Luca A, et al. The role of the EGFR signaling in tumor microenvironment. J Cell Physiol 214, 559–567 (2008).

31. Garcia-Palmero I, Villalobo A. Calmodulin regulates the translocation of Grb7 into the nucleus. FEBS letters 586, 1533–1539 (2012).

32. Tsai NP, Lin YL, Tsui YC, Wei LN. Dual action of epidermal growth factor: extracellular signal-stimulated nuclear-cytoplasmic export and coordinated translation of selected messenger RNA. J Cell Biol 188, 325–333 (2010).

33. Carpenter RL, Lo HW. STAT3 Target Genes Relevant to Human Cancers. Cancers (Basel) 6, 897–925 (2014).

34. Hachim IY, et al. Transforming Growth Factor-beta Regulation of Ephrin Type-A Receptor 4 Signaling in Breast Cancer Cellular Migration. Sci Rep 7, 14976 (2017).

35. Ding L, Shen Y, Ni J, Ou Y, Ou Y, Liu H. EphA4 promotes cell proliferation and cell adhesion-mediated drug resistance via the AKT pathway in multiple myeloma. Tumour Biol 39, 1010428317694298 (2017).

36. Saintigny P, et al. Global evaluation of Eph receptors and ephrins in lung adenocarcinomas identifies EphA4 as an inhibitor of cell migration and invasion. Molecular cancer therapeutics 11, 2021–2032 (2012).

37. Arpino G, Wiechmann L, Osborne CK, Schiff R. Crosstalk between the estrogen receptor and the HER tyrosine kinase receptor family: molecular mechanism and clinical implications for endocrine therapy resistance. Endocr Rev 29, 217–233 (2008).

38. Porter JC, Hogg N. Integrins take partners: cross-talk between integrins and other membrane receptors. Trends Cell Biol 8, 390–396 (1998).

39. Skog J, et al. Glioblastoma microvesicles transport RNA and proteins that promote tumour growth and provide diagnostic biomarkers. Nature cell biology 10, 1470–1476 (2008).

40. Chou SM, Huang TH, Chen HC, Li TK. Calcium-induced cleavage of DNA topoisomerase I involves the cytoplasmic-nuclear shuttling of calpain 2. Cellular and molecular life sciences: CMLS 68, 2769–2784 (2011).

41. Miao H, Nickel CH, Cantley LG, Bruggeman LA, Bennardo LN, Wang B. EphA kinase activation regulates HGF-induced epithelial branching morphogenesis. The Journal of cell biology 162, 1281–1292 (2003).

42. Gogas H, et al. Postoperative dose-dense sequential versus concomitant administration of epirubicin and paclitaxel in patients with node-positive breast cancer: 5-year results of the Hellenic Cooperative Oncology Group HE 10/00 phase III Trial. Breast cancer research and treatment 132, 609–619 (2012).

